# Human outbreak detection and best practice MPXV analysis and interpretation with squirrel

**DOI:** 10.1101/2025.08.13.669859

**Authors:** Áine O’Toole, Eddy Kinganda-Lusamaki, Rachel Colquhoun, Connor Chato, Emily Scher, Chris Kent, Sam Wilkinson, Josh Quick, Nick Loman, Ana T. Duggan, Placide Mbala-Kingebeni, Andrew Rambaut

**Affiliations:** Institute of Ecology & Evolution, University of Edinburgh, United Kingdom; Institut National de Recherche Biomédicale, Kinshasa, DR Congo; Service de Microbiologie, Département de Biologie Médicale, Cliniques Universitaires de Kinshasa, Université de Kinshasa, Kinshasa, Democratic Republic of the Congo; TransVIHMI, Université de Montpellier, INSERM, IRD, 34394 Montpellier, France; National Microbiology Laboratory, Public Health Agency of Canada, Winnipeg, Canada; Institute of Microbiology and Infection, University of Birmingham, United Kingdom; McMaster Ancient DNA Centre, Departments of Anthropology and Biochemistry, McMaster University, Hamilton, Canada

**Keywords:** MPXV, mpox, surveillance, software, phylogenetics, bioinformatics, APOBEC3, zoonosis, emerging disease, outbreak, genomic epidemiology

## Abstract

High numbers of reported mpox cases and recent identification of multiple sustained human outbreaks of MPXV have highlighted the need for robust, best-practice genomic surveillance tools. In light of the SARS-CoV-2 pandemic, many labs across the globe developed the capacity to do virus genome sequencing, however MPXV presents additional analytical challenges due to its large genome size, tracts of low-complexity or repeat regions, genetically distinct clades, and the need to perform bespoke APOBEC3-mutation reconstruction. We present squirrel (Some Quick Reconstruction to Resolve Evolutionary Links), an open source bioinformatic tool that can perform clade-aware alignment, mutation quality assessment, phylogenetic inference and automated APOBEC3-mutation classification on branches of the phylogeny. Squirrel can be run on the command line or launched through the EPI2ME GUI through the squirrel-nf workflow, enabling robust analysis without need for the command line. With the interactive output report produced and publication-ready APOBEC3-reconstruction visualisation, squirrel enables researchers to distinguish between zoonotic and sustained human outbreaks and help accurately inform public health responses.

## Introduction

MPXV, the virus that causes mpox, has historically been considered a zoonotic virus with limited capacity for sustained human-to-human transmission. However, our understanding of the epidemiology of mpox has shifted in recent years following a series of widespread, prolonged outbreaks in the human population. Whilst MPXV gained global attention in 2022 when it swept across the globe, evidence shows it had been spreading in the human population since at least 2016 (O’Toole et al. 2023). At time of writing, there are three ongoing sustained human outbreaks of MPXV alongside continued emergence of novel zoonotic cases of MPXV.

A key tool in understanding the dynamics of MPXV emergence has been the identification of APOBEC3-driven mutations as a signature for sustained transmission in the human population (Isidro et al. 2022; O’Toole et al. 2023; Gigante et al. 2022). This phenomenon enables researchers to infer whether the virus has been circulating in the human population and for how long, or whether the virus is likely caused by a recent zoonotic spillover. To answer these critical questions for new cases of mpox, public health labs and researchers have turned to genomic surveillance.

In the wake of the SARS-CoV-2 pandemic, many labs across the globe have developed the capacity for viral genomic sequencing, and this infrastructure and expertise can now be leveraged for MPXV genomic surveillance. The availability of ARTIC-style tiled amplicon sequencing approaches for MPXV (Kent et al. 2024; Chen et al. 2023) has facilitated broader uptake, as it can be readily adopted by those familiar with the SARS-CoV-2 sequencing protocol. However, sequencing and analysis of MPXV presents distinct challenges. MPXV is a large (∼200kb) double-stranded DNA virus with tracts of low-complexity and repetitive regions that can be problematic for some sequencing platforms. Moreover, MPXV has two genetically distinct clades and primer schemes optimized for one clade may perform poorly on the other. Mismatch between sequencing approach and analytical pipeline (e.g. metagenomic assembly pipeline for amplicon sequencing data) can result in low-quality genomes, incorrect SNP calls and incorporation of primer artefacts. These errors can distort phylogenetic analysis, alter the interpretation of the data and misinform public health responses.

In this manuscript, we present squirrel (Some Quick Reconstruction to Resolve Evolutionary Links), which was developed in order to provide robust, reliable genome alignment and phylogenetics for MPXV, as well as providing automated APOBEC3-mutation reconstruction across the reconstructed phylogeny, enabling more accurate and rapid analysis of MPXV genomes.

## Methods

### Availability and implementation

Squirrel is a command-line tool developed to provide best-practice analytic solutions for MPXV phylogenetics. It is Python-based with an embedded analysis pipeline built in a Snakemake framework (Köster and Rahmann 2018). All code is available on GitHub under an open-source GNU-GPL v3.0 licence at github.com/aineniamh/squirrel. Squirrel can be installed via bioconda, Docker or directly from the GitHub repository. For users not comfortable with the command line, squirrel can be run using Oxford Nanopore’s EPI2ME GUI platform (labs.epi2me.io) through the ARTIC Network resource squirrel-nf (https://github.com/artic-network/squirrel-nf). Documentation describing installation, the full usage and configuration options, and an example data file is available at github.com/aineniamh/squirrel.

### Pipeline description

Squirrel combines various quality control and phylogenetic functionality into one user-friendly software package. A user can submit a FASTA file of unaligned MPXV genome sequences to squirrel (Figure 2a). If the user specifies whether the ‘--clade’ is ‘cladei’ or ‘cladeii’, squirrel will select the respective RefSeq reference for either MPXV Clade I or II (Accession ID: NC_003310 or NC_063383). If the ‘--clade split’ flag is provided, squirrel will initiate the assignment pipeline that uses a custom MPXV random forest model to assign a clade to each input genome (Figure 2b). This model is an adapted version of the pangolin lineage assignment random forest model, pangoLEARN (O’Toole et al. 2021). The inputs are then split by major clade (Clade I or II) and a separate alignment pipeline is initiated for each clade detected in the input. Squirrel can also run on variola virus (VARV) and if ‘variola’ is specified the VARV RefSeq (Accession ID: NC_001611) is selected for alignment.

**Figure 1.**
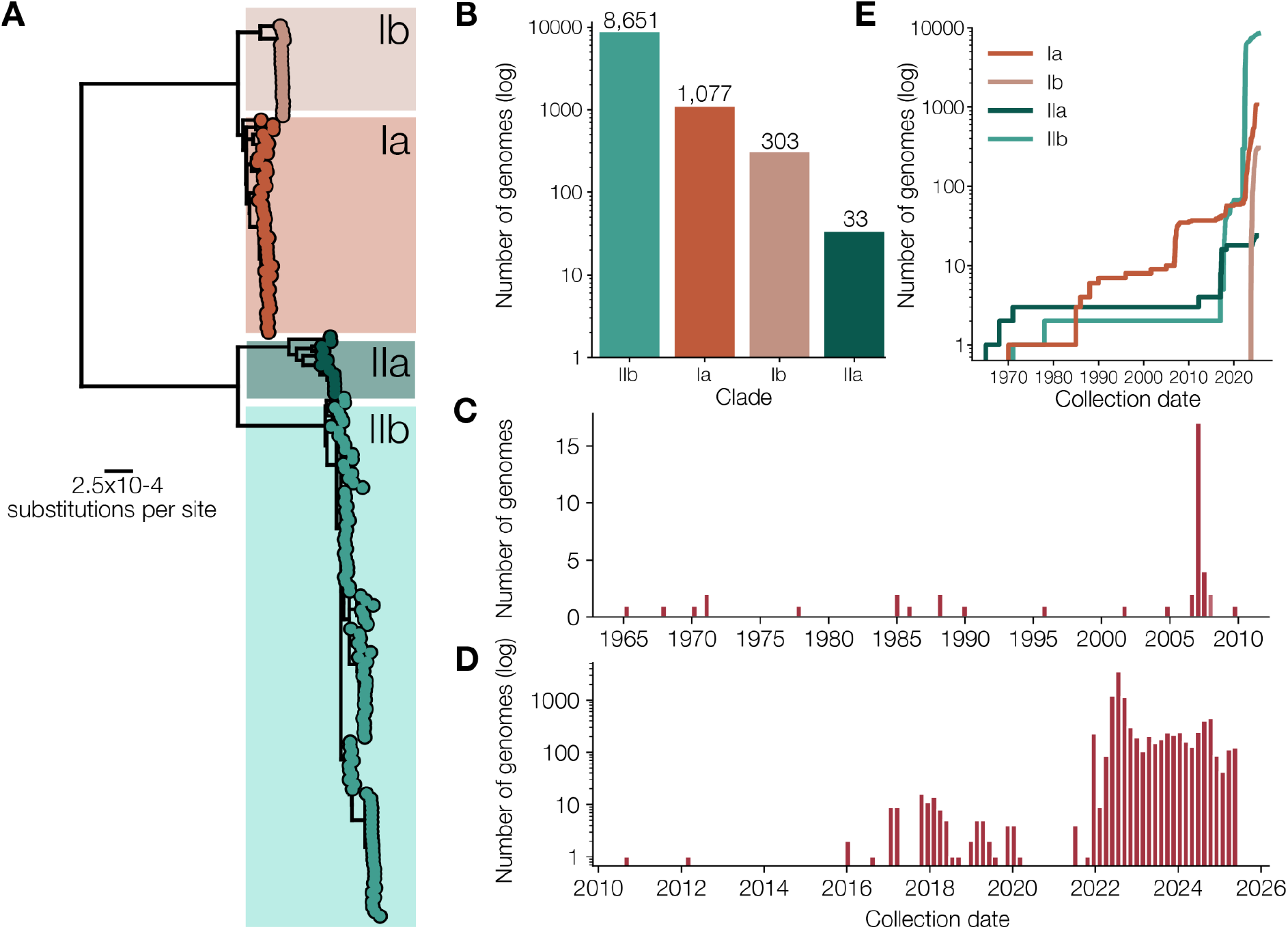
A) The diversity of MPXV is currently split into four clades: Clades Ia, Ib, IIa and IIb. B) Clade IIb has the most genomes with 8,651 sequences available on Pathoplexus, and Clades Ia, Ib and IIa have 1,077, 303 and 33 genomes respectively. C) Small numbers of samples were sequenced in the decades following MPXV discovery in 1958. D) Since the Clade IIb outbreak in 2017, many more genome sequences of the virus have been generated and shared online, with a total of 10,064 genomes on Pathoplexus that are sufficiently complete to assign clade. E) Clade Ia has been sampled and sequenced over the last few decades, however Clade Ib is recently discovered with the emergence of the sustained human outbreak sh2023, that emerged from Sud Kivu province in the Democratic Republic of Congo.

**Figure 2.**
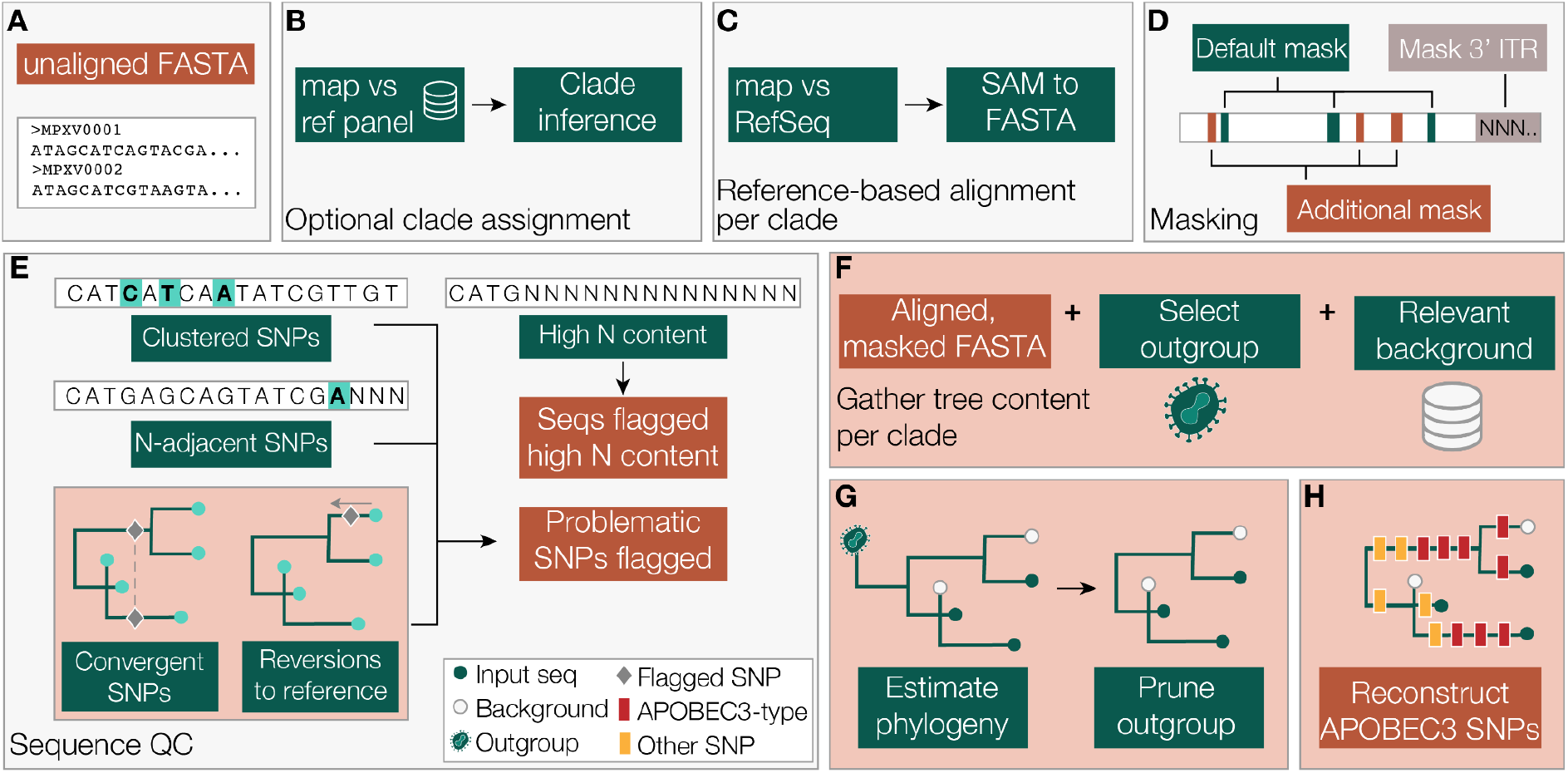
A) The user can input an unaligned multi-FASTA file of MPXV genomes. B) If ‘clade’ is set to ‘split’, the input genome sequences are assigned an MPXV clade using a custom random forest model trained on all MPXV data available on Pathoplexus. Sequences will then be split by major clade for any downstream analysis. C) The input sequences are mapped against the respective clade RefSeq genome and the output mapping file is converted to a FASTA alignment. D) Squirrel runs two kinds of masking on the input sequences. Firstly, the 3’ inverted terminal repeat (ITR) region is converted to N bases. A set of problematic sites are also masked, which are known low-complexity or repetitive regions that many of the known sequencing issues surround. This mask file is relatively minimal and users may wish to supply an additional mask file to supplement the masking on genomes. E) If squirrel is run in QC mode, it runs a module that generates a report flagging possibly problematic SNPs. SNPs flagged include clusters of SNPs, SNPs directly adjacent to an N, and also include convergent SNPs or SNPs that are reversions to the reference. This report is in the format that it can be used as the additional mask file in panel D. This file should be used to inform SNP investigation and confidence, and should not be used as a mask file without inspecting the SNPs themselves. F) Squirrel combines the input sequences with a clade-specific outgroup and optional background file. G) IQ-TREE 2 is used to estimate a maximum likelihood tree (with ancestral state reconstruction if APOBEC3-reconstruction mode is active). H) Squirrel reconstructs which mutations occur along which branch in the phylogeny and calculates whether they are consistent with APOBEC3-editing or not. It produces an output report describing each mutation and the context around it, such as gene, codon position, amino acid mutation, dinucleotide context. Squirrel produces an interactive HTML output report with the results summarised.

The alignment pipeline uses minimap2 to map all input sequences against the selected RefSeq reference genome and then gofasta (Jackson 2022) converts the output SAM file into a FASTA alignment (Figure 2c). The 3’ inverted terminal repeat region is masked out using Ns from the end of the genome (position 190,788 for MPXV or 184,722 for VARV) (Figure 2d). A set of known repetitive and low-complexity sites that are problematic for sequencing and alignment are also masked out with Ns. The full set of sites is hosted on the squirrel GitHub at github.com/aineniamh/squirrel/blob/main/squirrel/data. Additional sites can be provided to squirrel with the ‘--additional-mask’ flag and those nucleotide positions specified will also be masked from the alignment across all sequences (Figure 2d). Sites in specific sequences in the input FASTA file can be masked out by providing a ‘--sequence-mask’ csv file and problematic sequences can be entirely skipped from the input file by providing an ‘--exclude’ csv file. The masking function is run by default but both the ITR masking and the additional site masking can be toggled off.

When run with the optional ‘--seq-qc’ flag, squirrel will flag mutations that are directly adjacent to an N or a gap, a cluster of 3 or more mutations that occur within a 10 base window in the genome, and sequences with N content greater than 20%. Clusters of many SNPs in a small region of the genome can often be due to assembly or alignment errors. In a nucleotide sequence, N is often used to mask low-coverage regions, and a SNP directly adjacent to an N may be problematic. Similarly, a SNP next to a gap can be indicative of alignment issues and may require visual inspection of the alignment at that region. The user can then examine their data, decide whether the mutations or sequences are indeed problematic through visual inspection, and can then run a second round of squirrel and supply the output QC files as ‘--additional-mask’ or ‘--exclude’ files in order to construct an alignment with these additional flagged mutations masked out (Figure 2e).

Optionally, the user can use squirrel to do a maximum likelihood phylogenetic analysis, including APOBEC3 mutation reconstruction on the phylogeny. Based on the clade specified or assigned, squirrel will select an appropriate outgroup to add to the alignment (described in Table 1) to help correctly root the tree (Figure 2f). For ease of analysis of new data, squirrel has the option to include a separate background file in addition to the input FASTA file. This allows the user to have a maintained reference file that can be combined with new data for analysis in the report. Squirrel also contains a minimal reference set that can be automatically added if it is run with the ‘--include-background’ flag. We use IQ-TREE 2 (Minh et al. 2020) to build a maximum likelihood tree with a HKY substitution model (Hasegawa, Kishino, and Yano 1985) and later prune the outgroup from the resulting phylogeny using jclusterfunk (https://github.com/rambaut/jclusterfunk) (Figure 2g). If APOBEC3 phylogenetics is selected, IQ-TREE 2 is used to reconstruct the ancestral genomes on the internal nodes of the tree. Squirrel uses these state reconstruction to map mutations to branches in the phylogeny, assess whether they are consistent with APOBEC3 editing or not, calculate their position within coding frames and categorise the mutations by whether they are synonymous, non-synonymous, intergenic or nonsense mutations (Figure 2h).

**Table 1.**
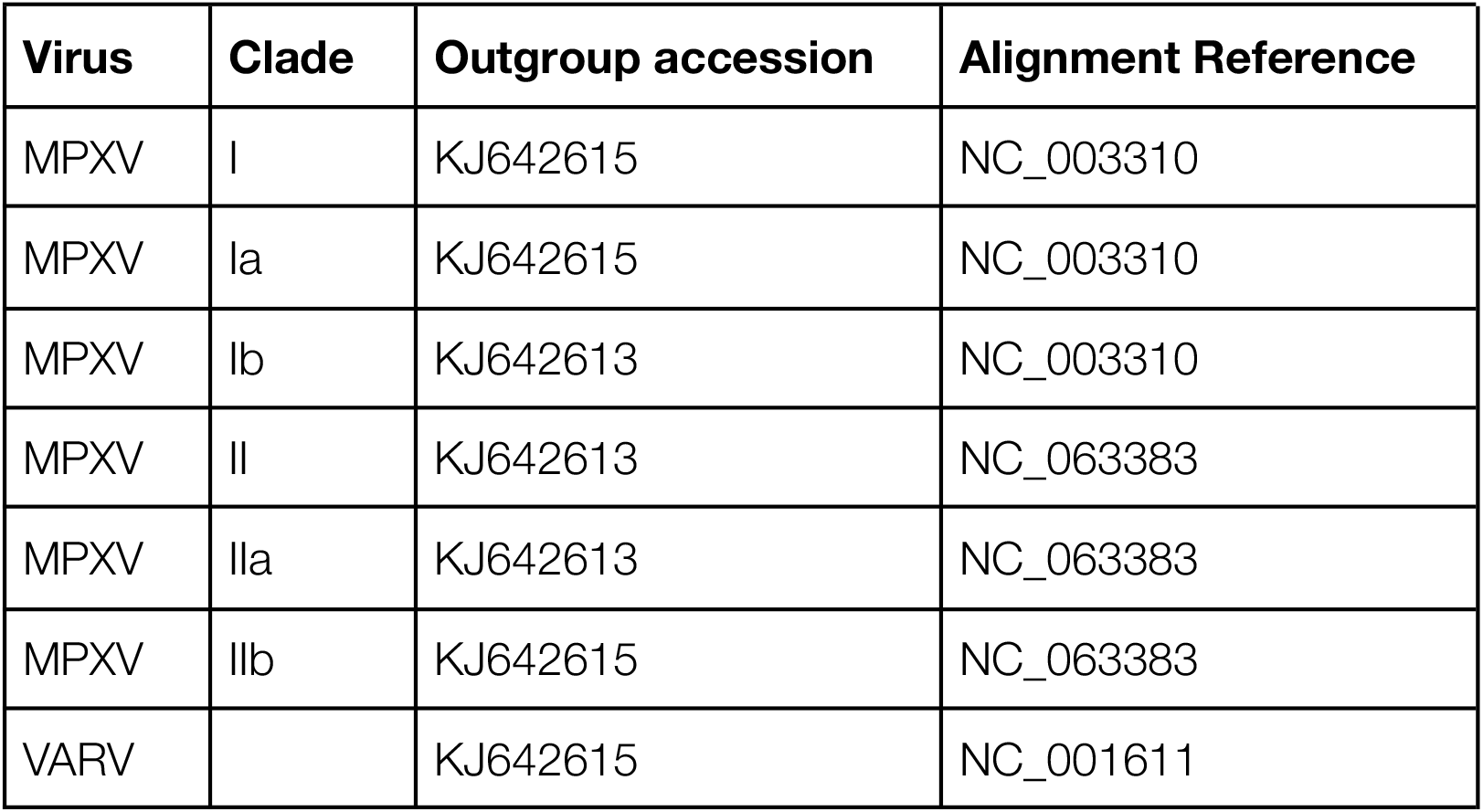
If not otherwise specified, squirrel will select an appropriate outgroup for phylogenetic analysis based on the clade or virus specified.

We use the phylogeny and reconstructed mutations to create a graphical representation of the phylogeny with APOBEC3 mutations annotated onto the branches (Figure 2h, Figure 3) using baltic (https://github.com/evogytis/baltic) and matplotlib. Mutations can be represented as coloured circles above the branches, or rectangles transecting the branches. APOBEC3-like mutations are shown in red and non-APOBEC3 mutations are shown in yellow. Squirrel can also produce an interactive tree figure with ape, ggtree and plotly using the ‘--interactive-tree’ flag (Yu 2020; Paradis and Schliep 2019).

**Figure 3.**
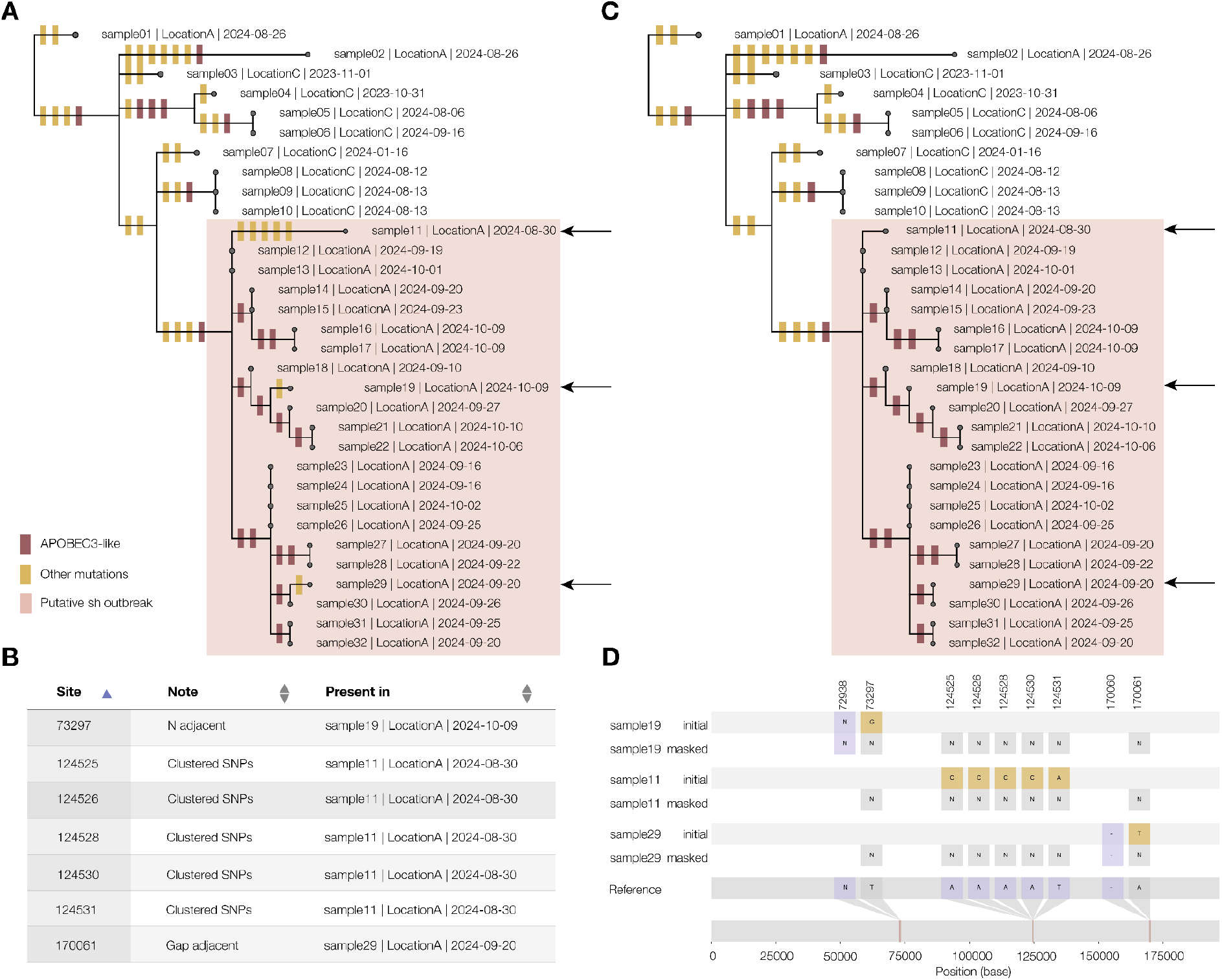
A) The squirrel output report displays the phylogeny of input genome sequences with reconstructed mutations along branches coloured by whether they are APOBEC3-like (red) or not (yellow). In this example, a subset of the sequences appear to have an excess of APOBEC3-like mutations along the branches, shaded and labelled manually here as a putative sustained human (sh) outbreak. Outside this clade, the branches show mutations more in line with the expected proportion of APOBEC3-mutations (∼12%). Within the putative outbreak clade, some sequences have non-APOBEC3 mutations (indicated by arrows), which is possible but would change our interpretation of the data. B) Running squirrel in QC mode produces an output table, displayed in the output report of sites that contain SNPs that may be due to bioinformatic error. SNPs are reported that lie directly adjacent to a gap or an N, and also if a high number of SNPs cluster together. C) Running squirrel again with the suggested mask file produces a report with the flagged SNPs masked out of the alignment. In this example case, the non-APOBEC3 mutations from the putative outbreak clade are masked out. D) With the additional mask, each flagged column in the alignment is replaced with N bases.

### Clade assignment model

We accessed all publicly available MPXV genome data with clade annotations on Pathoplexus.org and excluded any genomes that were <95% complete. The dataset included 7,622 Clade IIb, 363 Clade Ia, 98 Clade Ib and 33 Clade IIa genomes. We aligned all genomes against the Clade II RefSeq genome (Accession ID NC_063383) using squirrel in QC mode, and subsequently re-aligned the genomes with the additional mask file produced by squirrel, which identified 5,761 possibly problematic sites across the 8,116 genomes. We adapted the pangoLEARN random forest model to accept clade annotations and trained a new random forest model on the cleaned 8,116 genome dataset with clade annotations.

### Clade assignment tests

We compared three clade assignment methods for use within squirrel. Firstly, we used the command line Nextclade v3.15.3 with the mpoxall-clades dataset (version 2025-04-25--12-24-24Z), which is the standard clade assigner for many labs going MPXV genomics (Aksamentov et al. 2021). We also developed a pipeline whereby minimap2 v2.28-r1209 (Li 2018) is run (asm20) against a panel of curated background sequences representing each clade with only top hit recorded, and using the clade of the top hit as a clade assignment. Finally, we trained a custom random forest model on all available Pathoplexus MPXV data. This model was based on the pangoLEARN model used in pangolin for SARS-CoV-2 lineages (O’Toole et al. 2021). In order to assess the various methods of clade assignment, we selected a representative background set of genomes from each clade (Accession IDs in Table 2). For each of the genomes in this dataset, we first tested how the approaches behave with increasing ambiguity by simulating a dataset of 31,000 genomes with incrementing levels of random ambiguity (N-bases) from 5% to 95%. We then simulated genomes with incrementing genetic divergence to limit test the assignment models. To do this, we selected sites at random from across the MPXV genome, at increasing increments between 1 and 15% of sites, and *in silico* mutated them away from the original nucleotide at that site. We repeated this random selection 100 times to generate the test dataset. We created an extended dataset by further simulating genomes with 15-45% of sites randomly mutated in the same way.

**Table 2.** Sequences from which data was simulated to test the assignment models. *This record has since been removed from Genbank.

| Accession ID | Clade |
| --- | --- |
| MT903346.1 | Clade IIa |
| MT724769.1 | Clade IIa |
| AY741551.1 | Clade IIa |
| KP849470.1 | Clade IIa |
| DQ011156.1 | Clade IIa |
| KJ136820.1 | Clade IIa |
| MN346698.1 | Clade IIa |
| KJ642615.1 | Clade IIb |
| KJ642617.1 | Clade IIb |
| OP413718.1 | Clade IIb |
| MK783027.1 | Clade IIb |
| MK783033.1 | Clade IIb |
| MN648051.1 | Clade IIb |
| ON563414.3 | Clade IIb |
| OP415257.1 | Clade IIb |
| PP601220.1 | Clade Ib |
| PP601212.1 | Clade Ib |
| KJ642612.1 | Clade Ia |
| KJ642619.1 | Clade Ia |
| JX878412.1 | Clade Ia |
| JX878413.1 | Clade Ia |
| DQ011155.1 | Clade Ia |
| MT724770.1* | Clade Ia |
| KJ642613.1 | Clade Ia |
| PP601206.1 | Clade Ia |
| JX878423.1 | Clade Ia |
| DQ011154.1 | Clade Ia |

## Results

### Squirrel output report and QC functionality helps ensure data inference is robust

The output report produced by squirrel includes links to the alignment and tree file, and also a graphical representation of the phylogeny with APOBEC3-mutation reconstruction displayed on the branches (Figure 3a). If run in QC mode, the output report will contain a table listing the suggested sites to mask in another run, and the rationale behind the masking (Figure 3b). It is recommended to view these mutations in an alignment viewer (such as Geneious) to assess whether to mask these sites. Running squirrel again with the additional mask file will produce an alignment and phylogeny with the suggested columns masked out, which can help with interpretation of the data (Figure 3c-d).

### Random forest model clade assignment is robust to ambiguity and diversity

Clade assignment within squirrel was introduced in order to allow users to input data from a sequencing run with multiple clades, without needing the user to manually split the sequence file by clade, which could result in user error. We assessed a newly trained random forest model based on pangoLEARN, a custom pipeline involving minimap2 mapping against a reference panel and Nextclade parsimony placement on a mutation annotated tree. We tested the three methods against a dataset representing all four MPXV clades, with simulated ambiguity from 0% to 95% (n=31,000). We found that pangoLEARN was most robust to ambiguity in the data, in comparison to minimap2 and Nextclade, which did not assign a clade beyond 25% ambiguity (Figure 4). We also assessed the assignment approaches against a dataset of increasing divergence from known MPXV diversity by simulating mutations on a set of representative genomes from 1% to 15% divergence from original sequence (n=4,960). We found that pangoLEARN and Nextclade were both robust to increasing simulated increasing diversity beyond known diversity, however Nextclade required large memory requirements as the number of mutations increased (Figure 5). We simulated an additional dataset up to 45% divergence from known diversity, and even to this extreme distance pangoLEARN manages to accurately assign 70.3% of sequences (Supplementary Figure 1). The assignment approach using minimap2 was the least robust to data divergence from known diversity. For our purposes, we wanted a lightweight clade assignment method that was robust to both ambiguity and novel mutations and, based on our tests, the pangoLEARN random forest model was the most suitable.

**Figure 4.**
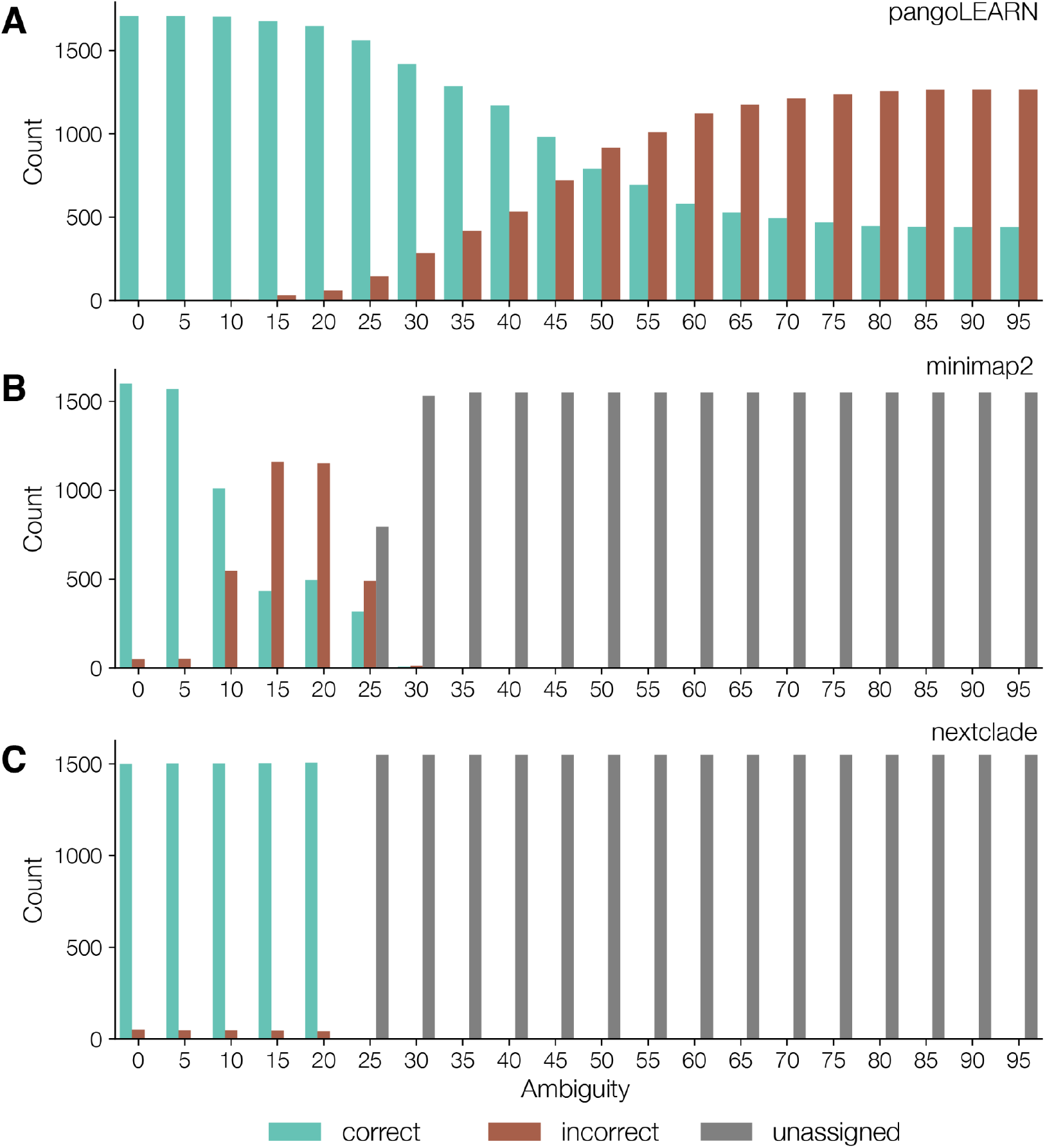
A) We evaluated the MPXV random forest clade assignment model on simulated sequences with increasing ambiguity (n=31,000). The model is robust to ambiguity and genomes that were 90% complete or more were all correctly assigned a clade. The model’s ability to correctly assign a clade declines as the ambiguity increases and as such clade assignments for very incomplete genomes should be interpreted with caution. B) As described in the minimap2 manual, it struggles to map beyond 20%, and the top hit becomes unreliable for clade assignment at greater than 10% ambiguity. C) We tested the command line version of Nextclade against the simulated ambiguous data and the model cuts off at 25% ambiguity, but is highly accurate to that point.

**Figure 5.**
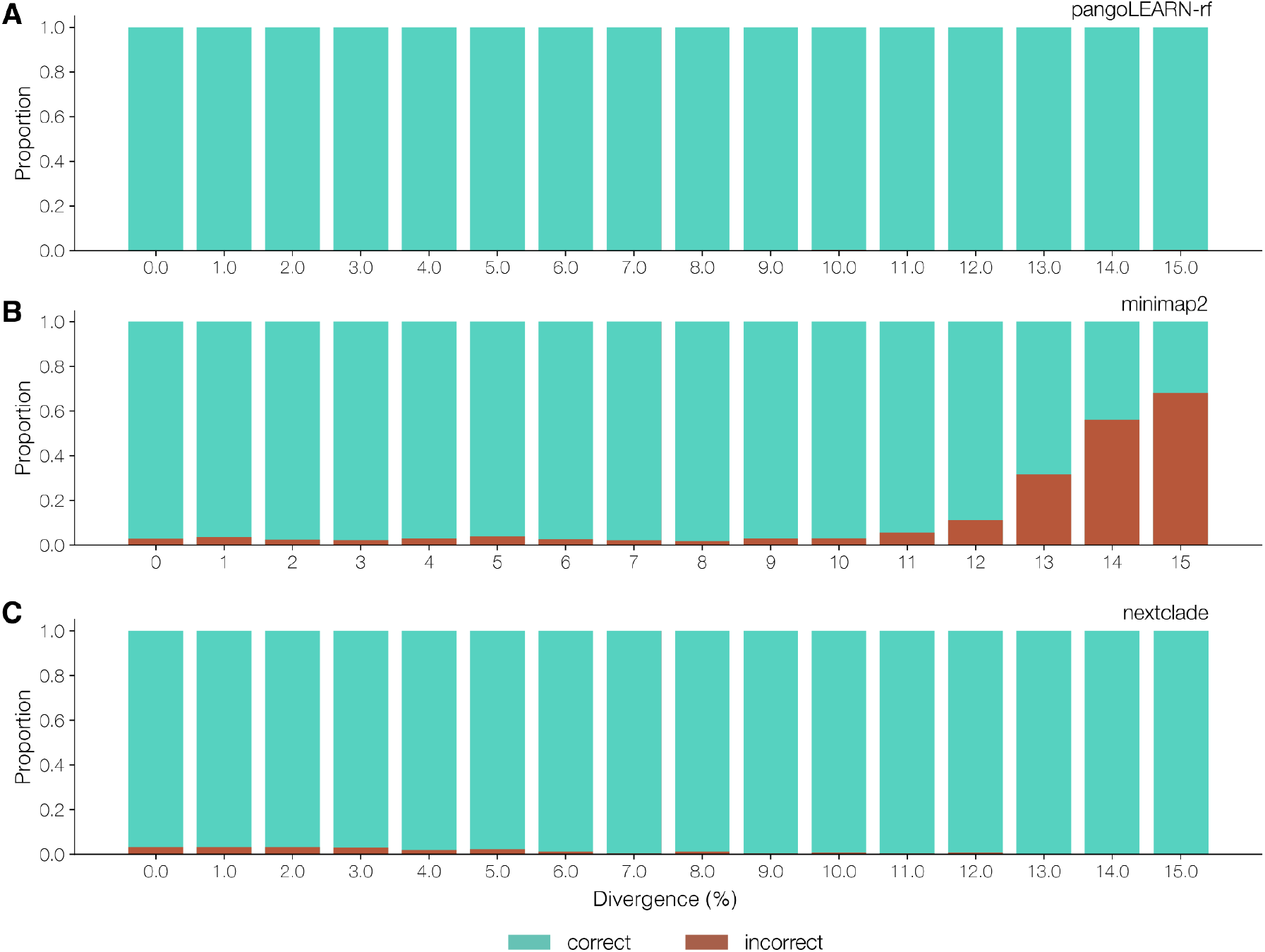
A) We assessed the performance of the MPXV pangoLEARN random forest model against a simulated dataset of increasing divergence from known diversity, based on representative sequences from each of the four MPXV clades (n=4,960). The pangoLEARN model was robust to divergence which is far beyond what we would observe in the MPXV population.

## Discussion

The recent surge of MPXV cases, particularly those involving sustained human-to-human transmission, has underscored a need for reliable genomic surveillance tools capable of distinguishing zoonotic spillovers from sustained human outbreaks. Whilst the increased global capacity for genomic surveillance following the SARS-CoV-2 pandemic has facilitated broader adoption of genomic epidemiology for MPXV, inconsistent genome data quality has highlighted a need for a reliable analytical tool tailored to MPXV to help troubleshoot mutation calls, generate robust alignments and perform APOBEC3-inference.

Squirrel was developed to meet this need by providing an end-to-end solution for MPXV genomic alignment, phylogenetic inference, mutation quality control, and APOBEC3 mutation reconstruction and classification. Squirrel supports both Clade I and Clade II alignments as well as analysis of variola virus. It can assign an MPXV clade to input genomes with its custom random forest clade assignment model, and will perform analysis on each clade with appropriately selected outgroups. Squirrel has been designed to be an adaptable tool, allowing the user to incorporate a default background set for context and to provide a custom background set, making it as streamlined as possible to combine existing and newly generated genome data in an analysis. It is thus far the only tool available that can perform the APOBEC3 reconstruction, which is a critical analysis for informing public health measures around any nascent MPXV outbreak.

Squirrel addresses a key analytical gap in MPXV genomic surveillance and the combination of its flexibility and ease of use makes it a valuable resource for both research and public health applications in the ongoing effort to monitor and control MPXV.

---

**Supplementary Figure 1.**
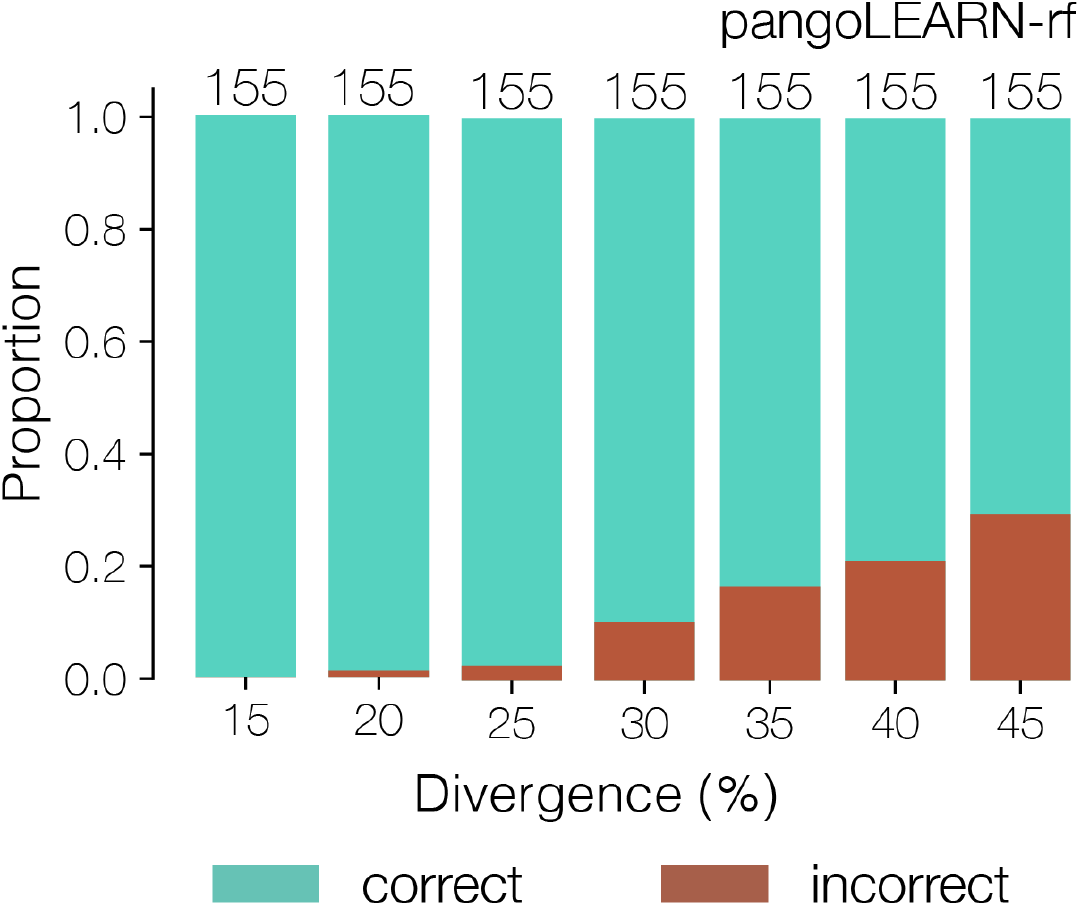
We simulated a dataset of up to 45% divergence from known MPXV diversity and found the pangoLEARN random forest model for MPXV clade assignment was robust to this diversity and still accurately assigned 70.3% of sequences at 45% divergence from known MPXV, which far exceeds known diversity within the species.

## Notes

### Competing Interest Statement

The authors have declared no competing interest.

### Funding

- Wellcome Trust: 313694/Z/24/Z

